# Salicylate, Bile Acids and Extreme Acid Cause Fitness Tradeoffs for Multidrug Pumps in *Escherichia coli* K-12

**DOI:** 10.1101/2020.11.21.392837

**Authors:** Samantha H. Schaffner, Abigail V. Lee, Minh T. N. Pham, Beimnet B. Kassaye, Haofan Li, Sheetal Tallada, Cassandra Lis, Mark Lang, Yangyang Liu, Nafeez Ahmed, Logan G. Galbraith, Jeremy P. Moore, Katarina M. Bischof, Joan L. Slonczewski

## Abstract

The aspirin derivative salicylate selects against bacterial multidrug efflux pumps of *Escherichia coli* K-12 such as MdtEF-TolC and EmrAB-TolC, and acid stress regulators such as GadE. Salicylate uptake is driven by the transmembrane pH gradient (ΔpH) and the proton motive force (PMF) which drives many efflux pumps. We used flow cytometry to measure the fitness tradeoffs of salicylate, bile acids, and extreme low pH for *E. coli* cultured with pump deletants. The AcrAB-TolC efflux pump conferred a fitness advantage in the presence of bile acids, an efflux substrate. Without bile acids, AcrA incurred a small fitness cost. The fitness advantage with bile acids was eliminated by the PMF uncoupler CCCP. The Gad acid fitness island encodes components of MdtEF-TolC (an acid-adapted efflux pump) as well as acid regulator GadE. The fitness advantage of *E. coli* cocultured with a Gad deletant (Δ*slp-gadX*) was lost in the presence of salicylate. Salicylate caused an even larger fitness cost for GadE. MdtE incurred negative or neutral fitness under all media conditions, as did EmrA. But when the competition cycle included two hours at pH 2, MdtE conferred a fitness advantage. The MdtE advantage required the presence of bile acids. Thus, the MdtEF-TolC pump is useful to *E. coli* for transient extreme acid exposure comparable to passage through the acidic stomach. Salicylate selects against some multidrug efflux pumps, whereas bile acids selects for them; and these fitness tradeoffs are amplified by extreme acid.

**IMPORTANCE:** Control of drug resistance in gut microbial communities is a compelling problem for human health. Growth of gut bacteria is limited by host-produced acids such as bile acids, and may be modulated by plant-derived acids such as salicylic acid. Membrane-soluble organic acids can control bacterial growth by disrupting membranes, decreasing cell pH, and depleting PMF. Our flow cytometry assay measures the fitness effects of exposure to membrane-soluble organic acids, with growth cycles that may include a period of extreme acid. We find that extreme-acid exposure leads to a fitness advantage for a multidrug pump, MdtEF-TolC, which otherwise incurs a large fitness cost. Thus, organic acids and stomach acid may play important roles in controlling multidrug resistance in the gut microbiome. Therapeutic acids might be developed to limit the prevalence of multidrug resistance pumps in environmental and host-associated communities.

## INTRODUCTION

Bacterial multi drug efflux systems (MDR) export diverse antibiotics, metals and harmful products of metabolism (1–3). These pumps serve as first-line defense against multiple antibiotics at low levels, while allowing time for the pathogen to acquire high-level resistance to a specific drug (4). Today, MDR pumps in pathogens pose a major threat to human health as pathogen multidrug resistance increases (5–7). Yet their evolution and persistence arise from their intrinsic contributions to bacterial survival that predate the antibiotic era. MDR pumps remove environmental or host-derived antimicrobials such as bile acids (8) as well as toxic products of the bacterium’s own metabolism (9,10).

The *E. coli* K-12 genome contains at least 36 multidrug efflux systems (9,11). The most active pump, AcrAB-TolC, exports antimicrobial drugs, antibiotics, dyes, organic solvents, essential oils, and hormones (12,13). AcrAB expression is upregulated by the global regulator MarA, which is activated by salicylate binding MarR repressor of MarRAB (14,15). Besides AcrAB-TolC, structurally similar pumps include the RND-type pump MdtEF-TolC (16,17) and Major Facilitator Superfamily (MFS) type pump EmrAB-TolC (18). The substrates of these pumps include bile acids such as cholic and deoxycholic acids (8,19).

The reversal of drug resistance, including that of MDR pumps, is a compelling goal (20). Control of MDR antibiotic resistance requires understanding of the physiological tradeoffs of maintaining such pumps. Physiological tradeoffs, including subtle changes in regulatory genes, can be explored by the tool of experimental evolution (21,22). Our laboratory uses experimental evolution to explore conditions that could reverse the relative fitness of drug efflux pumps (23–25). Aspirin derivatives such as benzoate and salicylate can select against resistance to antibiotics, favoring mutants that have lost multidrug efflux pumps. These acids also select against resistance to colistin (polymyxin E) in *Pseudomonas aeruginosa* (26).

A tradeoff of MDR pumps is that they require substantial amounts of energy (11,27). Some MDR pumps consume ATP, those of the ATP-binding cassette (ABC) family; but five major classes of MDR are driven by the electrochemical proton gradient, or proton motive force (PMF). For example, PMF drives pumps of the RND superfamily such as AcrAB-TolC (12,13); and the MFS superfamily such as EmrD (28). Multidrug efflux pumps use proton motive force (PMF) to expel a wide range of detrimental substances from the cell. However, these pumps waste energy when exposed to uncouplers of PMF from oxidative phosphorylation, such carbonyl cyanide m-chlorophenyl hydrazone (CCCP) (11,24). Full uncouplers transmit protons across the membrane both in protonated and unprotonated forms, whereas partial uncouplers transfer protons mainly in the protonated form. Partial uncouplers include membrane-soluble aromatic acids such as aspirin, salicylic acid, and ibuprofen, which belong to the class of nonsteroidal anti-inflammatory drugs (NSAIDs) (29,30). Many membrane-soluble acids and their derivatives are phytochemicals produced by plants that control their microbiomes in ways that are poorly understood.

Another common antimicrobial effect of organic acids is to destabilize the membrane. Gut microbes encounter membrane destabilization from bile acids, as well as cellular pH depression (18). The antimicrobial activity of bile acids largely structures the gut microbiome (31–33). Salicylate alters the structure of the outer membrane by regulating porins (34) and mediates oxidative stress (35) as well as pH depression. Membrane interactions are proposed to mediate the effect of salicylate on colistin resistance (26). Diverse organic acids may challenge bacteria in various ways, causing different fitness tradeoffs.

A less studied aspect of proton-driven drug pumps is their association with extreme-acid resistance (36), the ability of *E. coli* to survive transient exposure at a range of pH 1-3, as found in the human stomach (37,38). Moderate acid (pH 5-6.8) upregulates systems for extreme-acid survival such as those of the Gad acid fitness island (39,40). The acid fitness island includes genes for the MdtEF-TolC pump (17) and for Gad transcriptional regulators such as GadE (41). Gad-dependent acid resistance allows survival under extreme acid where *E. coli* cannot grow. The system raises cell pH by consuming protons through the decarboxylation of glutamate and glutamine by *gadA* and *gadB*, regulated by GadE. Despite the importance of this system, few studies of experimental evolution incorporate extreme-acid exposure; in one report, twenty daily cycles at pH 2.5 select for mutations in acid-resistance regulator EvgS (42).

We investigated the tradeoffs between PMF-driven MDR pumps and the presence of salicylate and bile acids (23–25). We developed a method using flow cytometry (43) to measure the relative fitness of MDR genes versus null alleles under various conditions, such as the presence of uncouplers and of exportable substrates such as bile acids (cholic acid and deoxycholic acid). We modified the method to include cycles of extreme acid exposure. Using this approach, we determined the conditions for relative fitness of AcrAB-TolC and EmrAB-TolC efflux pumps; and we showed that the MdtEF-TolC pump requires pH 2 exposure to confer positive relative fitness. This finding represents a novel case of an extreme acid-dependent drug efflux pump.

## RESULTS

### Relative fitness measurement by flow cytometry of YFP and CFP

We sought to measure the relative fitness of MDR genes in the presence of membrane-permeant acids. For this purpose we devised a flow cytometry assay modified from that of Wistrand-Yuen (Gullberg) (43) as described under Methods. In this assay, each strain had a gene of interest knocked out by replacement with *kanR* from alleles of the KEIO collection with the exception of the Δ*slp-gadX* strain (44). The *kanR* gene constitutively expresses an amino-glycoside 3’-phosphotransferase (44,45). In early competition trials, we found that *kanR* confers a small fitness cost compared to control strain *E. coli* W3110. Therefore, for all assays involving deletants expressing *kanR* our control strain included the allele*yhdN::kanR*. For the single gene deletions used in this experiment (Δ*acrA*::*kanR*, Δ*mdtE*::*kanR*, Δ*gadE*::*kanR*, and Δ*emrA*::*kanR*) each allele was transduced into a strain of *E. coli* K-12 W3110 containing the fluorophore alleles *galK*::*yfp* or *galK*::*cfp* inducible via a *lac* promoter (43).

The mixture of two strains was serially diluted 1000-fold each day, and observed over a total of 30 generations (doublings) from day zero to day 3. This generation number was sufficient to allow substantial fitness selection while minimizing the selection of secondary mutations (43). All culture media were buffered at pH 6.8, a level that allows cytoplasmic pH depression by membrane-permeant acids of lower pK_a_. Each day, a parallel dilution with IPTG inducer was performed using LBK-PIPES pH 6.8 buffered medium without stressors. The IPTG-induced populations express YFP or CFP for flow cytometry, whereas the overnight 1000-fold dilutions avoided energy-expensive fluorophore expression. This procedure enabled us to minimize the fitness effects of fluorophore expression during stress selection.

**Fig. 1A** shows the appearance of a distinctive population of YFP-expressing cells showing high fluorescence intensity with 488-nm excitation and low-intensity fluorescence with 405-nm excitation, versus a second population expressing CFP with low intensity at 488-nm excitation and high intensity with 405-nm excitation. **Fig. 1B** show a typical experiment in which the log_2_ ratios of YFP and CFP populations were reported for the tested mutant (W3110 *acrA*::*kanR*) and the control (W3110 *yhdN::kanR*). For each competition experiment, an equal number of trials were performed with the gene deletant strain expressing YFP and the control strain expressing CFP, and vice versa.

**Figure 1.**
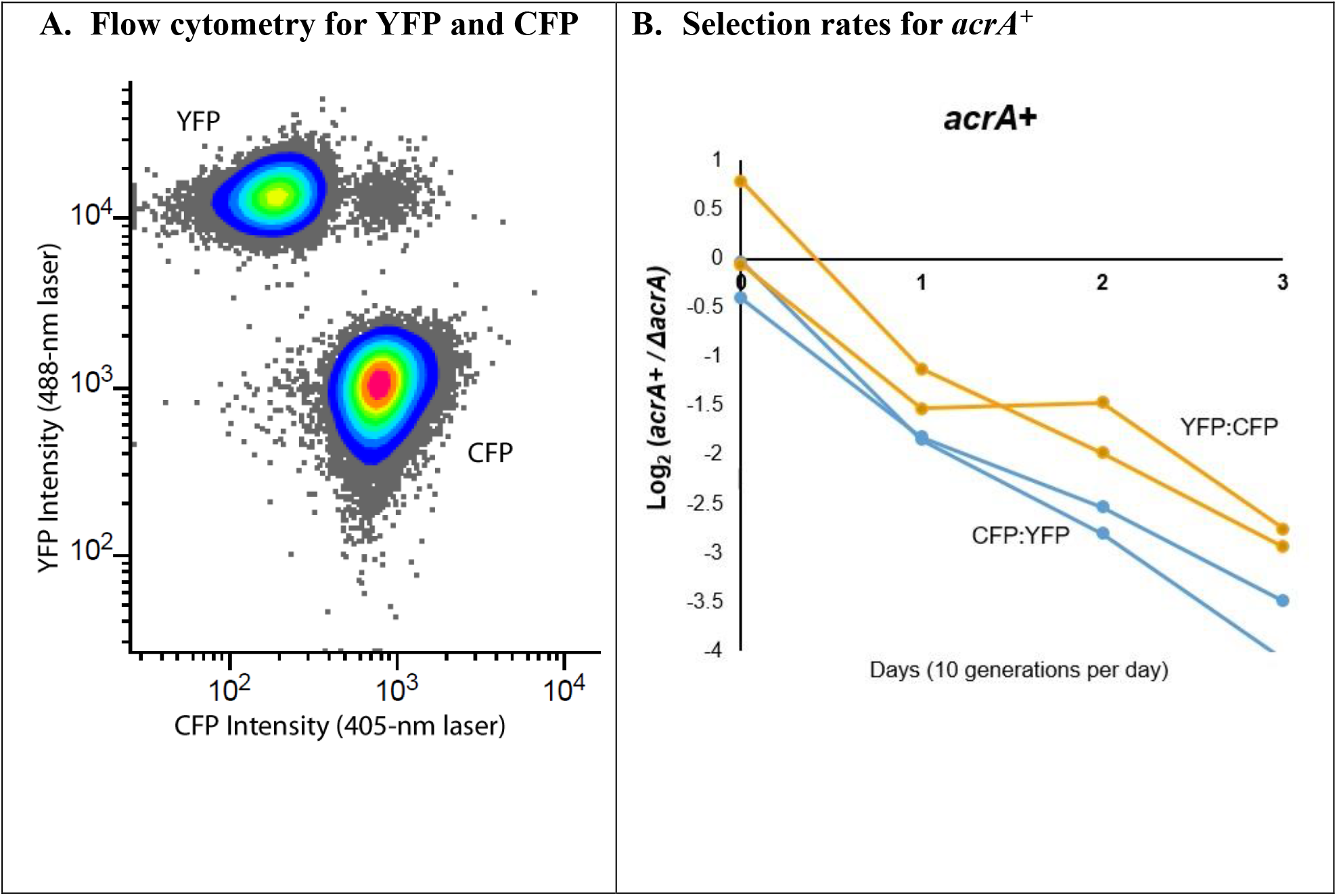
Flow cytometry competition assays of strains expressing YFP or CFP. **A.** Emission intensity indicates cells expressing cyan fluorescent protein (CFP) and yellow fluorescent protein (YFP) for two strains in coculture. Each culture is diluted and incubated for 2 h with IPTG to induce fluorophore expression. **B.** The slope of the log_2_ ratios for LBK-PIPES pH 6.8 control media (n=12) are compiled for Day 0 through Day 3 for all weeks tested. The absolute values of these ratios are taken from cell counts for W3110 Δ*yhdN::kanR* (*acrA^+^*) /W3110 *ΔacrA::kanR*. Each competition assay included an equal number of assays in which the *ΔyhdN* strain expressed YFP versus CFP. Cultures were diluted 1:1000 daily and assayed by flow cytometry as described under Methods. The slopes were calculated over days 0-3 of testing. The threshold for percent of each cell type is 0.01%.

Relative fitness was measured as the selection rate, the daily change in log_2_ ratio of cocultured strain populations (46). A selection rate of 1 unit per day indicates a two-fold increase in population of one strain compared to the competing strain’s population. This measure indicates the relative success of two strains cultured under a given stress condition.

### *AcrA* confers a fitness advantage in bile acids, but the advantage is eliminated by CCCP

AcrAB-TolC is the most widely studied of *E. coli* RND-type efflux pumps (47). A control strain expressing *acrA* was cocultured with the strain Δ*acrA*::*kanR* (**Fig. 2**). Cultured in pH 6.8 medium without stressors or in the presence of 6 mM salicylic acid, the *acrA^+^* strain showed a fitness deficit compared to Δ*acrA*::*kanR* (p<0.01, Fig. 2). By contrast, the presence of bile acids conferred a large advantage on the *acrA^+^* strain (p<0.001; Fig. 2). Thus, the AcrAB-TolC efflux pump is worth the energy cost when it is needed to export bile acids. Bile acids sustained a selective advantage for *acrA^+^* even in the presence of a low concentration of salicylate. The higher salicylate concentration (6 mM) with bile salts halted growth of both strains (data not shown).

**Figure 2.**
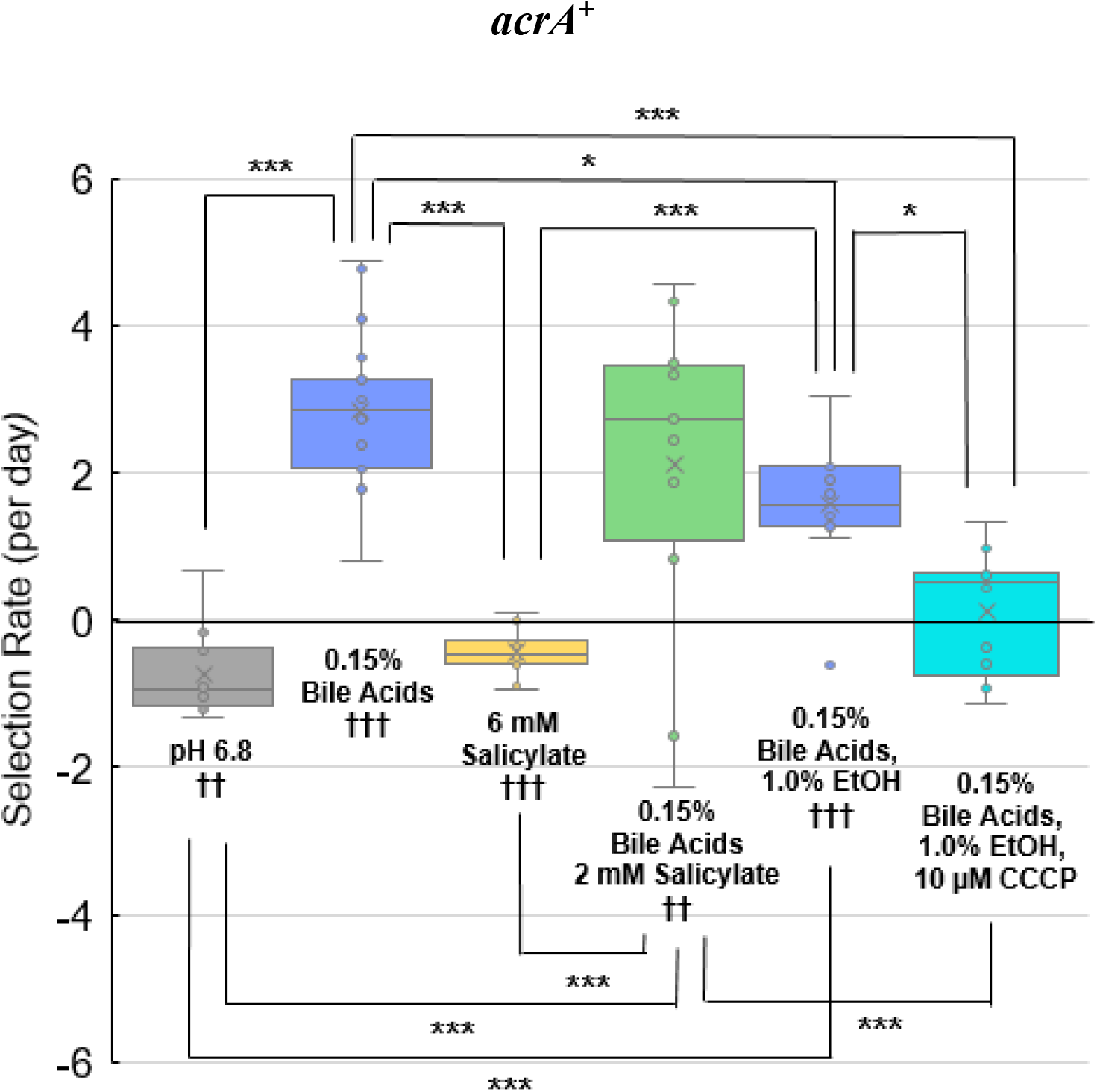
Selection for *acrA^+^* in the presence of salicylate, bile acids, and CCCP. Selection rate is given by log_2_(*ΔyhdN::kanR/ΔacrA::kanR*)/day. The control medium was LBK-PIPES pH 6.8 (n=12). Additional ingredients included 0.15% bile acids (n=20); 6 mM salicylate (n=12); 0.15% bile acids and 2 mM salicylate (n=12); 1.0% EtOH (n=12); and 10 μM CCCP (n=12). The slopes were calculated over days 0-3 of testing. For each condition, ANOVA and post-hoc Tukey tests were used to compare conditions to one another using brackets (*p<0.05, **p<0.01, ***p<0.001). Single sample t-tests were performed to compare each selection rate to a value of zero (†p<0.05, ††p<0.01, †††p<0.001).

Experimental evolution of *E. coli* in the presence of the PMF uncoupler CCCP selects for mutations in the AcrAB-TolC pump (24). We tested therefore whether CCCP would affect the relative fitness of *acrA^+^* in our assay **(Fig. 2)**. The fitness advantage of *acrA^+^* with bile salts was eliminated by the presence of 10 μM CCCP. The 1.0% EtOH control for CCCP stock solution showed little effect on the relative fitness of *acrA^+^* in 0.15% bile acids. Thus, depletion of PMF by CCCP eliminated the fitness advantage of AcrAB-TolC despite the presence of its key substrate for efflux.

### The Gad island confers a fitness advantage in bile acids that is neutralized by salicylate

The Gad acid fitness island includes genes whose products counteract acid stress such as periplasmic acid chaperones *hdeA* and *hdeB*, acid-resistance regulators *gadE* and *gadX*(39), and pump components *mdtEF* (48). Yet surprisingly, this region of the genome often undergoes deletion during experimental evolution in the presence of acid stressors (25,49).

Competition assays were conducted using a strain with most of the Gad island deleted by recombineering (Δ*slp-gadX*) (25). The parent strain W3310 had a fitness advantage over the Δ*slp-gadX* strain, in the presence or absence of bile acids (**Fig. 3A**). This advantage was reversed by the presence of 6mM sodium salicylate. Thus, the negative effect of salicylate overcomes whatever defense advantage is provided by overall components of the Gad island.

**Figure 3.**
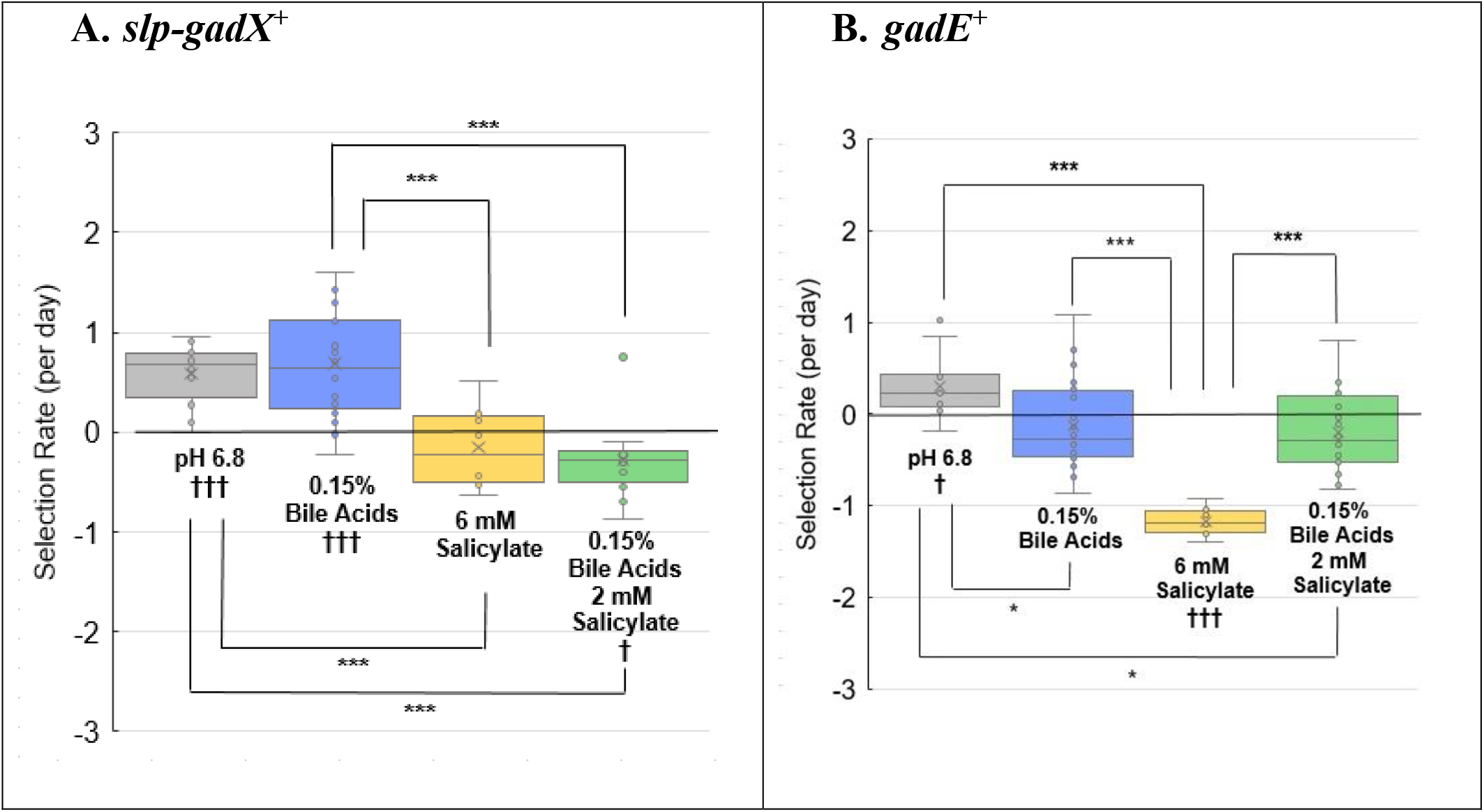
Selection for Gad island and for *gadE^+^*. **A.** Selection rate is given by log_2_(W3110 /*Δslp-gadX*) /day. Conditions were LBK-PIPES pH 6.8 (n=12) with 0.15% bile acids (n=20) or 6 mM salicylate (n=12), or 0.15% bile acids and 2 mM salicylate (n=12). Significance was determined as for Figure 2. **B.** Conditions include LBK-PIPES pH 6.8 (n=12) with 0.15% bile acids (n=20) or 6 mM salicylate (n=12), or 0.15% bile acids and 2 mM salicylate (n=16). Selection rate is given by log_2_(*ΔyhdN::kanR/ΔgadE::kanR*)/day. Significance was determined as for Figure 2.

### *GadE* incurs a fitness cost in the presence of salicylate

An important Gad component of the glutamate-dependent acid resistance pathway is *gadE* (50,51). GadE activates the decarboxylation of glutamate in the cell by the upregulation of *gadA* and *gadB* (52). Small adjustments of regulatory genes can affect drug resistance in unexpected ways (25,49). We sought to determine if salicylate affects the fitness advantage of *gadE^+^* similarly to that of the *slp-gadX* region overall.

In pH 6.8 medium, the control strain had a fitness advantage over *ΔgadE*, similar to the fitness advantage conferred by the Gad island as a whole (**Fig. 3B**). With bile acids, however, GadE conferred no fitness advantage. The presence of 6 mM salicylate, in the absence of bile acids, selected strongly against *gadE^+^* (p<0.001; **Fig. 3B**). These results suggest that some component regulated by GadE is sensitive to salicylate; and that some unidentified tradeoff exists with bile acids. We therefore tested the relative fitness conferred by *mdtE*, a GadE-regulated gene encoding a component of MdtEF-TolC.

### The fitness advantage of MdtE requires extreme acid exposure (pH 2)

The MdtEF-TolC efflux pump, when overexpressed in a strain lacking *acrB*, confers resistance to antibiotics and bile acids (53). The efflux complex is closely related to AcrAB-TolC; *mdtE* shares 55% homology with *acrA* (27) but shows unique acidic residues (17). We tested an *mdtE^+^* strain (W3110 *ΔyhdN::kanR*) against W3110 *ΔmdtE::kanR* in the presence of 6mM salicylate and 0.15% bile acids. The *mdtE^+^* strain incurred a fitness cost in the control medium buffered at pH 6.8, as well as with 0.15% bile acids, or with 6 mM salicylate (p<0.001; **Fig. 4A**). All conditions tested showed a fitness cost or neutral selection for MdtEF-TolC.

**Figure 4.**
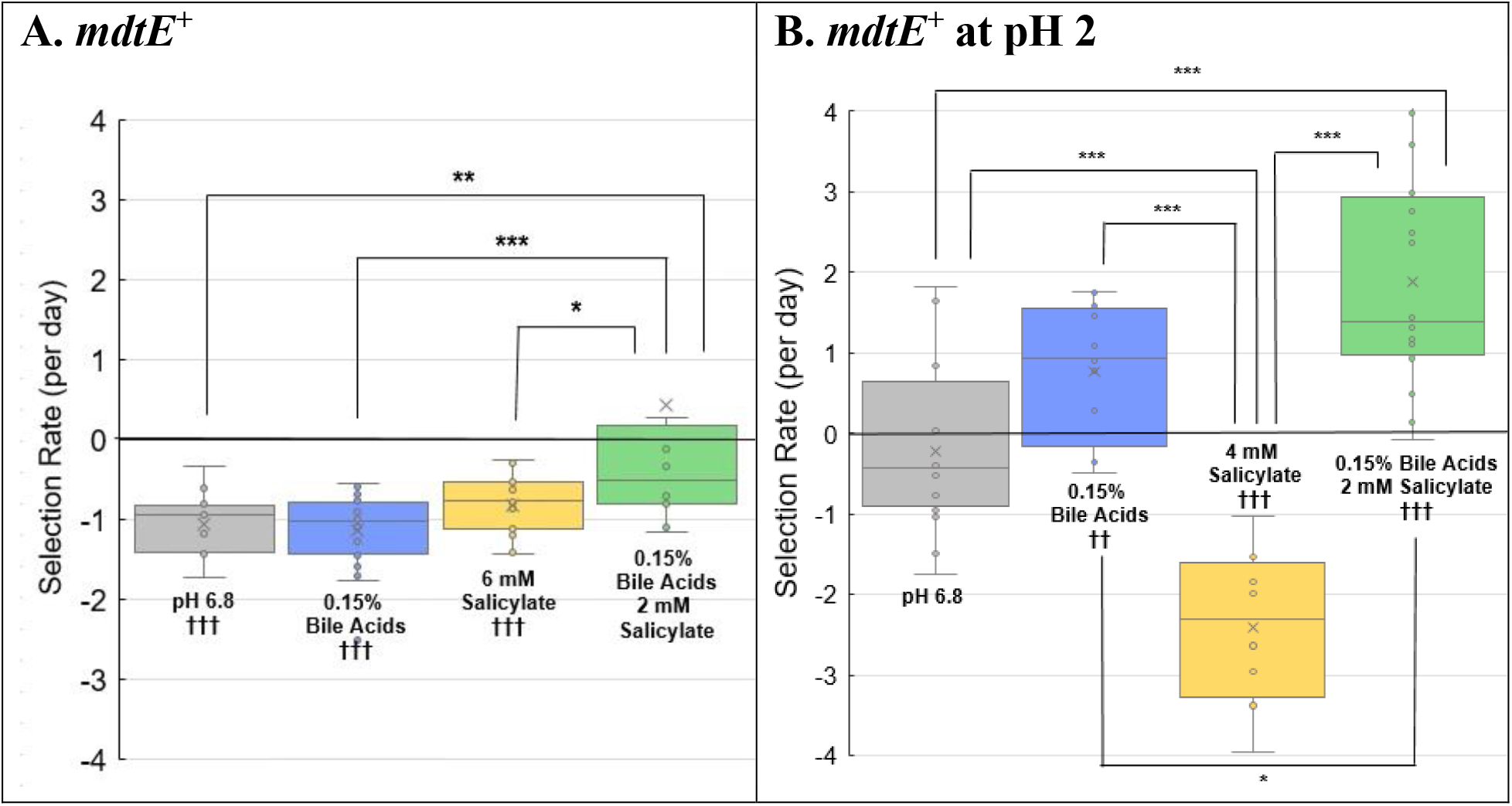
Selection for *mdtE^+^* with or without pH 2 exposure. Selection rate is given by log_2_(*ΔyhdN::kanR/ΔmdtE::kanR*)/day. Significance was determined as for Figure 2. **A**. LBK-PIPES pH 6.8 (n=16) with 0.15% bile acids (n=24); 6 mM salicylate (n=16); or 0.15% bile acids and 2 mM salicylate (n=12) **B**. LBK-PIPES pH 6.8 (n=16) with 0.15% bile acids (n=16); 4 mM salicylate (n=8); or 0.15% bile acids and 2 mM salicylate (n=16). The daily growth cycle included 100-fold dilution in unbuffered LBK pH 2, with incubation for 2 h followed by a 10-fold dilution in the appropriate competition media adjusted to pH 7.0, yielding a final pH of 6.8.

Under what conditions therefore does *mdtE* confer a fitness advantage, and thus maintain the pump’s persistence in the genome? The MdtEF-TolC efflux pump exports many substrates in common with AcrAB-TolC, but its distal pocket includes amino-acid residues with a lower isoelectric point (pI=3.1 for MdtEF, pI=4.0 for AcrAB) (17). Thus, the function of MdtEF may be optimized for extreme low pH, in common with other Gad components such as the periplasmic acid chaperones HdeA and HdeB.

So we tested the possibility that *mdeE* confers a fitness advantage during pH 2 exposure, a condition for which the Gad regulon maintains extreme-acid survival (36). This extreme-acid condition mimics the passage of bacteria through an acidic stomach, an adaptation that enteric *E. coli* must retain in the genome. Starting on Day zero, the CFP/YFP strain mixture was diluted 100-fold in unbuffered LBK at pH 2.0. The acidified cell suspension, approximately pH 2, was incubated for 2 hours. Under this condition, survival of *E. coli* K-12 strains is typically 50-100% (36,52,54). The acidified suspension was then diluted ten-fold in media buffered with PIPES at pH 7.0, leading to a final pH of 6.8. The total dilution overall was 1,000-fold per day, comparable to our original relative fitness assay. This period of pH 2 exposure was conducted for each of the three days of the assay.

The exposure to extreme acid eliminated the fitness cost of MdtE in media at pH 6.8, and led to a fitness advantage in the presence of bile acids, with or without 2 mM salicylate (**Fig. 4B**). In the higher concentration of 4 mM salicylate, however, MdtE incurred a large fitness cost (p<0.001). The higher salicylate concentration (6 mM) did not allow sustained growth. Note that after 100-fold dilution in LBK pH 2, the medium would still contain 0.06 mM salicylate. At external pH 2, where the *E. coli* internal pH is about pH 5 (54) the uptake of salicylic acid (with a pK_a_ = 2.8) would be greatly increased by the transmembrane pH difference, a ΔpH of approximately three units. By contrast, the pK_a_ values of cholic and deoxycholic acids are higher, 5.0 and 6.6 respectively, so the ΔpH would drive less uptake.

### EmrA confers a fitness loss in bile acids

The EmrAB-TolC efflux pump structurally resembles AcrAB-TolC efflux pump, with EmrA providing a link between the outer membrane efflux pump component, TolC and inner membrane component, EmrB (55). The EmrAB-TolC efflux pump is activated by salicylate, and confers resistance to related compounds as well as CCCP (18,56). As seen for *mdtE*, expression of *emrA* incurred a fitness cost in the pH 6.8, in the presence or absence of 0.15% bile acids (**Fig. 5**). Neutral fitness advantage was conferred in the presence of bile acids and salicylate.

**Figure 5.**
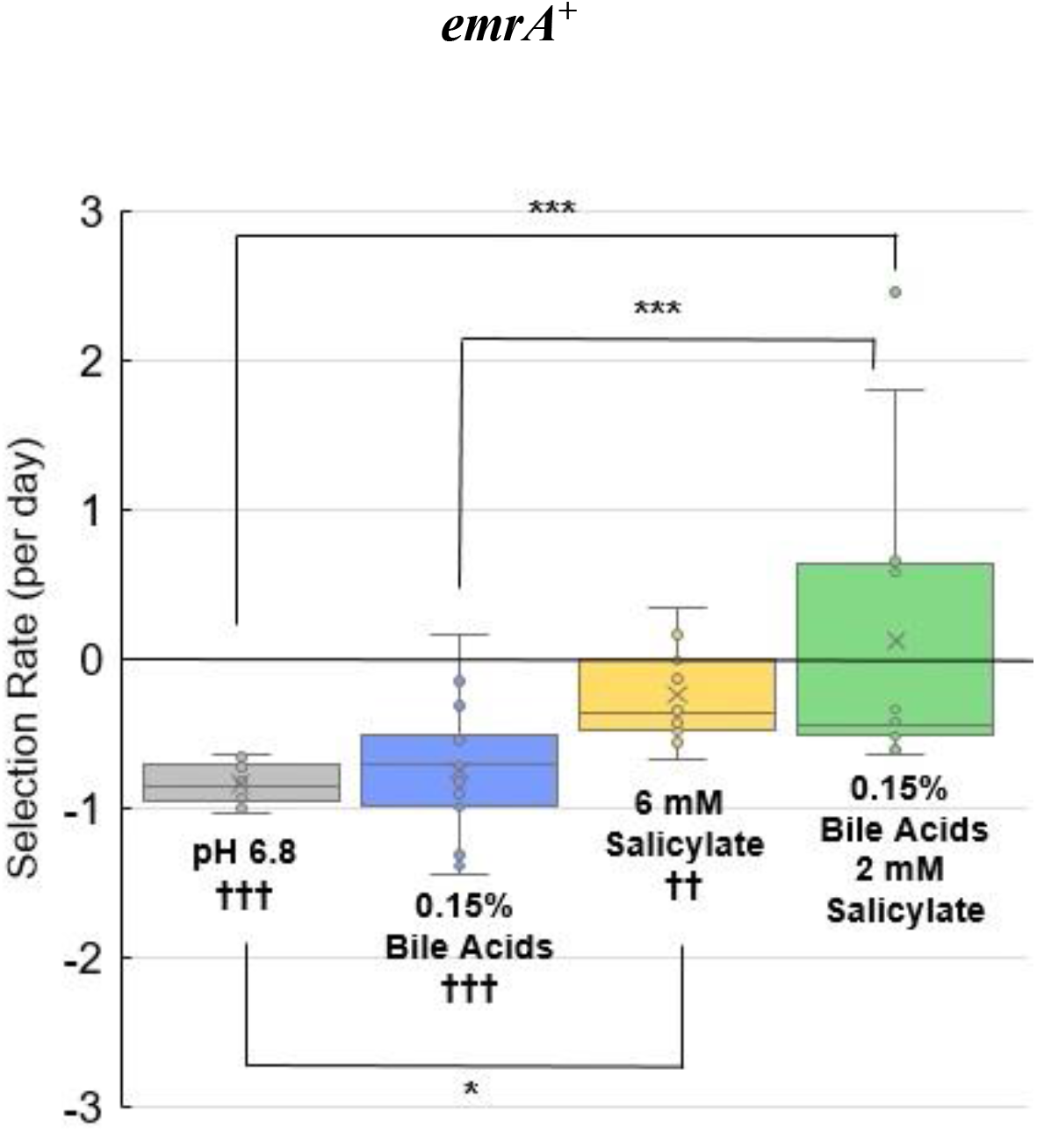
Selection for *emrA^+^*. Selection rate is given by log_2_(*ΔyhdN::kanR/ΔemrA::kanR*)/day. Media include LBK-PIPES pH 6.8 (n=16) with 0.15% bile acids (n=16); 6 mM salicylate (n=16); or 0.15% bile acids and 2 mM salicylate (n=12). Significance was determined as for Figure 2.

## DISCUSSION

Multidrug efflux pumps are critical for their ability to export a range of detrimental compounds and metabolites (1–3). However, these pumps consume PMF when not needed and may decrease relative fitness. Thus, the strain with wild-type AcrA incurs a fitness cost in control medium (**Fig. 2**) yet with 0.15% bile acids, the pump is strongly favored. The loss of the AcrA fitness advantage in the presence of CCCP is consistent with the selection for *ΔacrAB* mutants during experimental evolution with CCCP (24).

By contrast, under similar conditions of exposure to bile acids, the MdtEF-TolC and EmrAB-TolC pumps showed no selective advantage. This may be because AcrAB-TolC plays the dominant role for bile-acid efflux; with this pump functional, the other pumps are not needed. The selective value of possessing multiple multiple-substrate efflux pumps has not been clear; but it would make sense if their functions are optimized for diverse environmental contitions.The requirement of pH 2 exposure for positive fitness of the MdtEF-TolC efflux pump was unexpected (**Fig. 4**). It has not been known why the MdtEF pump is part of the Gad acid fitness island, along with a complex set of regulators of acid resistance via glutamate decarboxylase (57). Yet the fitness advantage with pH 2 exposure suggests that this pump effluxes bile acids under extreme acid. Bile acids are found in highest concentration in the lumen of the small intestine, where they aid lipid absorption (58–60). While bile does not normally reach the stomach, many patients exhibit chronic bile reflux gastritis (61) a condition associated with disorders such as carcinogenesis. Thus, pumps capable of extreme acid-dependent efflux of bile acids could be useful for bacteria that experience gastric transit. Just as there are “pH specialist” enzymes such as penicillin binding proteins (62) there may be pH specialist drug efflux pumps.

The effect of salicylate on the relative fitness of *gadE*, and of *mdtE*, is consistent with the surprising findings that experimental evolution in benzoate (23) or in moderate acid permitting growth (pH 4.8) selects for loss of the Gad island, as well as various acid-related regulators (49). Unless cells experience the pH 2 exposure, there is no long-term advantage to maintaining the MdtEF pump.

The overall Gad island (**Fig. 3A**) did confer a fitness advantage, except in the presence of salicylate. Part of the advantage may derive from the Gad genes other than *mdtEF*. The Gad regulon includes various components whose functions in acid resistance are poorly understood (25). The *hdeAB* acid stress operon encodes two periplasmic proteins that prevent protein aggregation at pH 2.0 (63,64). It may be more advantageous for the cell to lose one energetically expensive pump rather than their entire acid defense system.

Our modified flow cytometry assay enables us to explore the fitness effects of extreme-acid exposure comparable to that of the human stomach, which enteric pathogens must pass through to reach the intestinal tract (36,52,54). Our results may provide clues as to the relative fitness of drug-exporting strains during human uptake of aromatic acid medications such as aspirin. The mechanisms of antimicrobial action of aromatic acids may be more complex than uncoupling of PMF, including PMF-amplified uptake and membrane disruption (23,25,26,65). We will explore further the effect of acid exposure on fitness of EmrA and other MDR pumps, once the pandemic situation allows return of our student researchers to the lab.

The measurement of relative fitness under acid exposure has pharmaceutical implications (66). Gastric levels of salicylate or other bioactive agents, from the diet or or from drug therapy, could affect the ability of drug-resistant pathogens to enter the stomach and colonize the intestinal tract. Including cycles of extreme acid in testing may be useful for assessing oral pharmaceuticals.

## MATERIALS AND METHODS

### Strains and media

All strains used in our experiments are derived from *E. coli* K-12 W3110 (**Table 1**). Strains were constructed by phage P1 transduction (67). The main growth medium was LBK broth (10 g/l tryptone, 5 g/l yeast extract, 7.45 g/l potassium chloride) buffered at pH 6.8 with 100 mM piperazine-N,N’-bis(2-ethanesulfonic acid (PIPES, pK_a_= 6.80) using NaOH to adjust pH (25). This medium is designated LBK-PIPES pH 6.8. Sodium salicylate, bile acids (50/50 mixture of sodium cholate and sodium deoxycholate) and carbonyl cyanide m-chlorophenyl hydrazone (CCCP) were all obtained from Millipore Sigma.

**Table 1.**
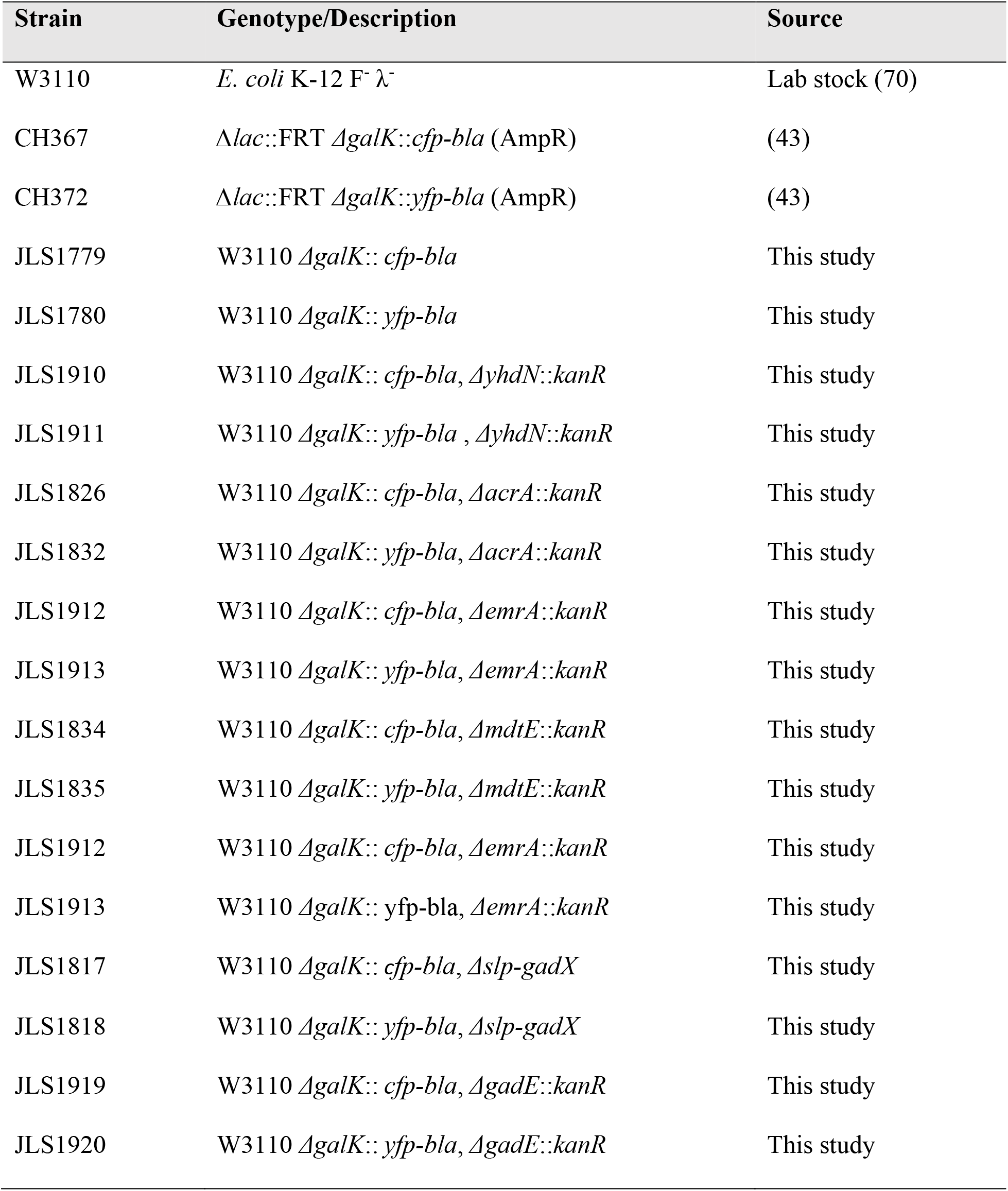
*E. coli* strains used in this study

### FACS Competition Assays

Relative fitness of cocultured strains was measured by flow cytometry of strains expressing yellow fluorescent protein (YFP) or cyan fluorescent protein (CFP) (43). Strains were cultured in 2 mL of LBK-PIPES pH 6.8 incubated in separate tubes at 37°C with rotation for 16 h. On the next day (Day-1) 20 μl of each inoculated culture was pipetted into 2 ml of the appropriate competition media for which the strains were competed and incubated for 24 h at 37°C with rotation. On Day 0, for each experimental condition 2 μl of a 1:1 mixture of the CFP and YFP strains to compete was added to 2 ml of the appropriate competition medium and incubated at 37°C with rotation for 24 h. From this culture, serial dilutions were repeated (2 μl into 2 ml) on Days 1 and 2.

To include pH 2 exposure, samples were incubated in unbuffered LBK pH 2 for the first 2 h of incubation each day (Day 0 through Day 2). Each day, 2 μl of the mixture of competing CFP and YFP strains was added to 200 μl of unbuffered LBK pH 2 and incubated at 37°C for 2 h. After this incubation period, 1.8 mL of the competition media buffered to pH 7 was added (restoring pH to approximately 6.8) and the tubes were incubated for 24 h at 37°C.

For daily flow cytometry: A separate dilution of each CFP and YFP coculture was performed by adding 50 μl (1:40 dilution) of the 1:1 mixtures of labeled strains and 20 μl (1:100 dilution) of 100 mM IPTG to 2 ml of LBK-PIPES pH 6.8. This tube was incubated for 2 h at 37 °C while rotating and then sampled using the BD FACSMelody Cell Sorter using a blue laser (488 nm) and violet laser (405 nm). A 545/20 filter was used for YFP emission, and a 528/45 filter for CFP emission. Each competition mixture was run for 50,000 total events and diluted with LBK-PIPES pH 6.8 so that the processed events were greater than 98% and the event rate was less than 10,000 events/second. The threshold value of counts for each tested strain in a run was set at 0.01%. Two technical replicates of each competition mixture were recorded and averaged. The percentages of cells with YFP or CFP fluorescence was recorded. This process was repeated each day for days 0-3 of testing. For each experimental condition, unless specified otherwise, 12 biological replicates were averaged, always with equal numbers of YFP to CFP and CFP to YFP competitions.

For each gene tested, the selection rate *s* was calculated as:

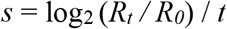

where *R* represents the ratio of cell numbers for the control strain (W3110 Δ*yhdN* or W3110) to the deletant strain of interest; and *t* represents time in days (with daily dilution 1:1,000, approximately 10 generations per day) (46,68,69). This rate gives a biological indication of the change in population distribution of cocultured genetic variants over time. For example, a selection rate of 1 unit per day (one two-fold increase per 10 generations) means that each day, one of two cocultured strains increases its population advantage two-fold over the competing strain.

### Statistical Analysis

Significance for selection rate differences was determined by ANOVA and post-hoc Tukey tests using R. In the figures, brackets indicate the Tukey results (*p<0.05, **p<0.01, ***p<0.001). Single sample t-tests were performed to compare the value of each selection rate to zero (†p<0.05, ††p<0.01, †††p<0.001). Supplemental File 1 provides all results of statistical calculations.

## ACKNOWLEDGMENTS

This work was supported by National Science Foundation awards MCB-1923077 and MRI-1725426. We thank Erik Wistrand-Yuen and Dan Andersson for valuable discussions and for the generous gift of strains.

## REFERENCES

1. Du D, Wang-Kan X, Neuberger A, van Veen HW, Pos KM, Piddock LJV, et al. Multidrug efflux pumps: structure, function and regulation. Nat Rev Microbiol [Internet]. 2018;16(9):523–39. Available from: http://dx.doi.org/10.1038/s41579-018-0048-6

2. Sulavik MC, Houseweart C, Cramer C, Jiwani N, Murgolo N, Greene J, et al. Antibiotic susceptibility profiles of Escherichia coli strains lacking multidrug efflux pump genes. Antimicrob Agents Chemother. 2001;45(4):1126–36.

3. Tikhonova EB, Zgurskaya HI. AcrA, AcrB, and TolC of Escherichia coli form a stable intermembrane multidrug efflux complex. J Biol Chem. 2004;279(31):32116–24.

4. Nolivos S, Cayron J, Dedieu A, Page A, Delolme F, Lesterlin C. Role of AcrAB-TolC multidrug efflux pump in drug-resistance acquisition by plasmid transfer. Science. 2019;364(6442):778–82.

5. Ventola CL. The antibiotic resistance crisis: causes and threats. P T J. 2015;40(4):277–83.

6. Malik B, Bhattacharyya S. Antibiotic drug-resistance as a complex system driven by socio-economic growth and antibiotic misuse. Sci Rep. 2019;9(9788):1–12.

7. Spengler G, Kincses A, Gajdács M, Amaral L. New roads leading to old destinations: Efflux pumps as targets to reverse multidrug resistance in bacteria. Molecules. 2017;22(3).

8. Thanassi DG, Cheng LW, Nikaido H. Active efflux of bile salts by Escherichia coli. J Bacteriol. 1997;179(8):2512–8.

9. Teelucksingh T, Thompson LK, Cox G. The Evolutionary Conservation of Escherichia coli Drug Efflux Pumps Supports Physiological Functions. J Bacteriol. 2020;(August):1–40.

10. Rosner JL, Martin RG. An excretory function for the Escherichia coli outer membrane pore TolC: Upregulation of marA and soxS transcription and Rob activity due to metabolites accumulated in tolC mutants. J Bacteriol. 2009;191(16):5283–92.

11. Paulsen IT, Brown MH, Skurray RA. Proton-dependent multidrug efflux systems. Microbiological Reviews. 1996.

12. Du D, Wang Z, James NR, Voss JE, Klimont E, Ohene-Agyei T, et al. Structure of the AcrAB-TolC multidrug efflux pump. Nature [Internet]. 2014;509(7501):512–5. Available from: http://dx.doi.org/10.1038/nature13205

13. Shi X, Chen M, Yu Z, Bell JM, Wang H, Forrester I, et al. In situ structure and assembly of the multidrug efflux pump AcrAB-TolC. Nat Commun [Internet]. 2019;10(1):4–9. Available from: http://dx.doi.org/10.1038/s41467-019-10512-6

14. Cohen SP, Levy SB, Foulds J, Rosner JL. Salicylate induction of antibiotic resistance in *Escherichia coli*: Activation of the *mar* operon and a *mar*-independent pathway. J Bacteriol. 1993;175(24):7856–62.

15. Wang T, Kunze C, Dunlop MJ. Salicylate Increases Fitness Cost Associated with MarA-Mediated Antibiotic Resistance. Biophys J [Internet]. 2019;117(3):563–71. Available from: https://doi.org/10.1016/j.bpj.2019.07.005

16. Nishino K, Nikaido E, Yamaguchi A. Regulation and physiological function of multidrug efflux pumps in Escherichia coli and Salmonella. Biochim Biophys Acta - Proteins Proteomics [Internet]. 2009;1794(5):834–43. Available from: http://dx.doi.org/10.1016/j.bbapap.2009.02.002

17. Novoa D, Otakuye C-B. The Anaerobic Efflux Pump MdtEF-TolC Confers Resistance to Cationic Biocides. bioRXiv Prepr http://dx.doi.org/101101/036806. 2019;

18. Lomovskaya O, Lewis K. emr, an Escherichia coli locus for multidrug resistance. Proc Natl Acad Sci U S A. 1992;

19. Pos KM. Drug transport mechanism of the AcrB efflux pump. Biochim Biophys Acta - Proteins Proteomics [Internet]. 2009;1794(5):782–93. Available from: http://dx.doi.org/10.1016/j.bbapap.2008.12.015

20. Andersson DI, Hughes D. Antibiotic resistance and its cost: is it possible to reverse resistance ? Nat Publ Gr [Internet]. 2010;8(4):260–71. Available from: http://dx.doi.org/10.1038/nrmicro2319

21. Blount ZD, Barrick JE, Davidson CJ, Lenski RE. Genomic analysis of a key innovation in an experimental *Escherichia coli* population. Nature [Internet]. 2012;489(7417):513–8. Available from: http://dx.doi.org/10.1038/nature11514 http://www.nature.com/nature/journal/v489/n7417/pdf/nature11514.pdf

22. Blount ZD, Maddamsetti R, Grant NA, Ahmed ST, Jagdish T, Baxter JA, et al. Genomic and phenotypic evolution of escherichia coli in a novel citrate-only resource environment. Elife. 2020;9:1–64.

23. Creamer KE, Ditmars FS, Basting PJ, Kunka KS, Hamdallah IN, Bush SP, et al. Benzoate-and salicylate-tolerant strains of *Escherichia coli* K-12 lose antibiotic resistance during laboratory evolution. Appl Environ Microbiol. 2017;83(2):e02736.

24. Griffith JM, Basting PJ, Bischof KM, Wrona EP, Kunka KS, Tancredi AC, et al. Experimental evolution of Escherichia coli K-12 in the presence of proton motive force (PMF) uncoupler carbonyl cyanide m-chlorophenylhydrazone selects for mutations affecting PMF-driven drug efflux pumps. Appl Environ Microbiol. 2018;

25. Moore JP, Li H, Engmann ML, Bischof KM, Kunka KS, Harris ME, et al. Inverted Regulation of Multidrug Efflux Pumps, Acid Resistance, and Porins in Benzoate-Evolved Escherichia coli K-12. Appl Environ Microbiol. 2019;85(16):1–21.

26. Malla CF, Mireles NA, Ramírez AS, Poveda JB, Tavío MM. Aspirin, sodium benzoate and sodium salicylate reverse resistance to colistin in Enterobacteriaceae and Pseudomonas aeruginosa. J Antimicrob Chemother. 2020;

27. Anes J, McCusker MP, Fanning S, Martins M. The ins and outs of RND efflux pumps in Escherichia coli. Front Microbiol. 2015;6(JUN):1–14.

28. Lee J, Sands ZA, Biggin PC. A numbering system for MFS transporter proteins. Front Mol Biosci. 2016;3(JUN):1–13.

29. Cardinale DA, Lilja M, Mandic M, Gustafsson T, Larsen FJ, Lundberg TR. Resistance training with co-ingestion of anti-inflammatory drugs attenuates mitochondrial function. Front Physiol. 2017;

30. Mahmud T, Rafi SS, Scott DL, Wrigglesworth JM, Bjarnason I. Nonsteroidal antiinflammatory drugs and uncoupling of mitochondrial oxidative phosphorylation. Arthritis Rheum. 1996;

31. Martins A, Spengler G, Rodrigues L, Viveiros M, Ramos J, Martins M, et al. pH modulation of efflux pump activity of multi-drug resistant Escherichia coli: Protection during its passage and eventual colonization of the colon. PLoS One. 2009;4(8).

32. Schubert K, Olde Damink SWM, von Bergen M, Schaap FG. Interactions between bile salts, gut microbiota, and hepatic innate immunity. Immunol Rev. 2017;279(1):23–35.

33. Kurdi P, Kawanishi K, Mizutani K, Yokota A. Mechanism of growth inhibition by free bile acids in lactobacilli and bifidobacteria. J Bacteriol. 2006;

34. Rosner JL, Chai TJ, Foulds J. Regulation of OmpF porin expression by salicylate in Escherichia coli. J Bacteriol. 1991;173(18):5631–8.

35. Pomposiello PJ, Bennik MHJ, Demple B. Genome-wide transcriptional profiling of the Escherichia coli responses to superoxide stress and sodium salicylate. J Bacteriol. 2001;183(13):3890–902.

36. Foster JW. *Escherichia coli* acid resistance: Tales of an amateur acidophile. Nat Rev Microbiol [Internet]. 2004 Nov [cited 2015 Oct 25];2:898–907. Available from: http://www.ncbi.nlm.nih.gov/pubmed/15494746

37. Dressman JB, Berardi RR, Dermentzoglou LC, Russell TL, Schmaltz SP, Barnett JL, et al. Upper Gastrointestinal (GI) pH in Young, Healthy Men and Women. Pharm Res An Off J Am Assoc Pharm Sci. 1990;

38. Beasley DE, Koltz AM, Lambert JE, Fierer N, Dunn RR. The evolution of stomach acidity and its relevance to the human microbiome. PLoS One. 2015;

39. Mates AK, Sayed AK, Foster JW. Products of the *Escherichia coli* acid fitness island attenuate metabolite stress at extremely low pH and mediate a cell density-dependent acid resistance. J Bacteriol. 2007;189(7):2759–68.

40. Deng Z, Shan Y, Pan Q, Gao X, Yan A. Anaerobic expression of the gadE-mdtEF multidrug efflux operon is primarily regulated by the two-component system ArcBA through antagonizing the H-NS mediated repression. Front Microbiol. 2013;4(JUL):1–14.

41. Ma Z, Richard H, Tucker DL, Conway T, Foster JW. Collaborative regulation of Escherichia coli glutamate-dependent acid resistance by two AraC-like regulators, GadX and GadW (YhiW). J Bacteriol. 2002;

42. Johnson MD, Bell J, Clarke K, Chandler R, Pathak P, Xia Y, et al. Characterization of mutations in the PAS domain of the EvgS sensor kinase selected by laboratory evolution for acid resistance in Escherichia coli. Mol Microbiol. 2014;93(5):911–27.

43. Gullberg E, Cao S, Berg OG, Ilbäck C, Sandegren L, Hughes D, et al. Selection of resistant bacteria at very low antibiotic concentrations. PLoS Pathog. 2011;7(7):e1002158.

44. Baba T, Ara T, Hasegawa M, Takai Y, Okumura Y, Baba M, et al. Construction of *Escherichia coli* K-12 in-frame, single-gene knockout mutants: The Keio collection. Mol Syst Biol [Internet]. 2006;2(1):0008. Available from: http://www.pubmedcentral.nih.gov/articlerender.fcgi?artid=1681482&tool=pmcentrez&rendertype=abstract

45. Datsenko KA, Wanner BL. One-step inactivation of chromosomal genes in Escherichia coli K-12 using PCR products. Proc Natl Acad Sci U S A. 2000;97(12):6640–5.

46. Burghardt LT, Epstein B, Guhlin J, Nelson MS, Taylor MR, Young ND, et al. Select and resequence reveals relative fitness of bacteria in symbiotic and free-living environments. Proc Natl Acad Sci U S A. 2018;115(10):2425–30.

47. Anes J, McCusker MP, Fanning S, Martins M. The ins and outs of RND efflux pumps in Escherichia coli. Front Microbiol. 2015;6(JUN):1–14.

48. Nishino K, Senda Y, Yamaguchi A. The AraC-family regulator GadX enhances multidrug resistance in *Escherichia coli* by activating expression of *mdtEF* multidrug efflux genes. J Infect Chemother. 2008;14(1):23–9.

49. He A, Penix SR, Basting PJ, Griffith JM, Creamer KE, Camperchioli D, et al. Acid evolution of *Escherichia coli* K-12 eliminates amino acid decarboxylases and reregulates catabolism. Appl Environ Microbiol. 2017;83:e00442–17.

50. Ma Z, Gong S, Richard H, Tucker DL, Conway T, Foster JW. GadE (YhiE) activates glutamate decarboxylase-dependent acid resistance in *Escherichia coli* K-12. Mol Microbiol. 2003;49(5):1309–20.

51. Castanié-Cornet MP, Cam K, Bastiat B, Cros A, Bordes P, Gutierrez C. Acid stress response in Escherichia coli: Mechanism of regulation of gadA transcription by RcsB and GadE. Nucleic Acids Res. 2010;38(11):3546–54.

52. Richard H, Foster JW. Escherichia coli glutamate-and arginine-dependent acid resistance systems increase internal pH and reverse transmembrane potential. J Bacteriol. 2004;186(18):6032–41.

53. Nishino K, Senda Y, Yamaguchi A. The AraC-family regulator GadX enhances multidrug resistance in Escherichia coli by activating expression of mdtEF multidrug efflux genes. J Infect Chemother. 2008;14(1):23–9.

54. Small P, Blankenhorn D, Welty D, Zinser E, Slonczewski JL. Acid and base resistance in *Escherichia coli* and *Shigella flexneri:* role of *rpoS* and growth pH. J Bacteriol [Internet]. 1994 Mar [cited 2016 Aug 8];176(6):1729–37. Available from: http://www.ncbi.nlm.nih.gov/pubmed/8132468

55. Borges-Walmsley MI, Beauchamp J, Kelly SM, Jumel K, Candlish D, Harding SE, et al. Identification of oligomerization and drug-binding domains of the membrane fusion protein EmrA. J Biol Chem. 2003;278(15):12903–12.

56. Lomovskaya O, Lewis K, Matin A. EmrR is a negative regulator of the Escherichia coli multidrug resistance pump emrAB. J Bacteriol. 1995;177(9):2328–34.

57. Waterman SR, Small PLC. Transcriptional expression of Escherichia coli glutamate-dependent acid resistance genes gadA and gadBC in an hns rpoS mutant. J Bacteriol. 2003;185(15):4644–7.

58. Hofmann AF. The continuing importance of bile acids in liver and intestinal disease. Archives of Internal Medicine. 1999.

59. Hamilton JP, Xie G, Raufman JP, Hogan S, Griffin TL, Packard CA, et al. Human cecal bile acids: Concentration and spectrum. Am J Physiol - Gastrointest Liver Physiol. 2007;293(1):256–63.

60. Hegyi P, Maléth J, Walters JR, Hofmann AF, Keely SJ. Guts and gall: Bile acids in regulation of intestinal epithelial function in health and disease. Physiol Rev. 2018;98(4):1983–2023.

61. Vere CC, Cazacu S, Comanescu V, Mogoanta L, Rogoveanu I, Ciurea T. Endoscopical and histological features in bile reflux gastritis. Rom J Morphol Embryol. 2005;46(4):269–74.

62. Mueller EA, Egan AJF, Breukink E, Vollmer W, Levin PA. Plasticity of Escherichia coli cell wall metabolism promotes fitness and antibiotic resistance across environmental conditions. Elife. 2019;8:1–24.

63. Kern R, Malki A, Abdallah J, Tagourti J, Richarme G. Escherichia coli HdeB is an acid stress chaperone. J Bacteriol. 2007;

64. Hong W, Jiao W, Hu J, Zhang J, Liu C, Fu X, et al. Periplasmic protein HdeA exhibits chaperone-like activity exclusively within stomach pH range by transforming into disordered conformation. J Biol Chem. 2005;280(29):27029–34.

65. Sundaramoorthy NS, Suresh P, Selva Ganesan S, GaneshPrasad AK, Nagarajan S. Restoring colistin sensitivity in colistin-resistant E. coli: Combinatorial use of MarR inhibitor with efflux pump inhibitor. Sci Rep [Internet]. 2019;9(1):1–13. Available from: http://dx.doi.org/10.1038/s41598-019-56325-x

66. Abuhelwa AY, Williams DB, Upton RN, Foster DJR. Food, gastrointestinal pH, and models of oral drug absorption. Eur J Pharm Biopharm [Internet]. 2017;112:234–48. Available from: http://dx.doi.org/10.1016/j.ejpb.2016.11.034

67. Deininger KNW, Horikawa A, Kitko RD, Tatsumi R, Rosner JL, Wachi M, et al. A requirement of tolc and MDR efflux pumps for acid adaptation and GadAB induction in escherichia coli. PLoS One. 2011;

68. Dykhuizen DE. Experimental Studies of Natural Selection in Bacteria. Annu Rev Ecol Syst. 1990;21(1):373–98.

69. Lenski RE, Rose MR, Simpson SC, Tadler SC. Long-term experimental evolution in *Escherichia coli*. I. Adaptation and divergence during 2,000 generations. Am Nat. 1991;138:1315–41.

70. Hayashi K, Morooka N, Yamamoto Y, Fujita K, Isono K, Choi S, et al. Highly accurate genome sequences of Escherichia coli K-12 strains MG1655 and W3110. Mol Syst Biol [Internet]. 2006 [cited 2016 Aug 8];2:2006.0007. Available from: http://www.ncbi.nlm.nih.gov/pubmed/16738553

